# Modeling the relative role of human mobility, land-use and climate factors on dengue outbreak emergence in Sri Lanka

**DOI:** 10.1101/462150

**Authors:** Ying Zhang, Jefferson Riera, Kayla Ostrow, Sauleh Siddiqui, Harendra de Silva, Sahotra Sarkar, Lakkumar Fernando, Lauren Gardner

## Abstract

**Background:** More than 80,000 dengue cases including 215 deaths were reported nationally in less than seven months between 2016-2017, a fourfold increase in the number of reported cases compared to the average number over 2010-2016. The region of Negombo, located in the Western province, experienced the greatest number of dengue cases in the country and is the focus area of our study, where we aim to capture the spatial-temporal dynamics of dengue transmission.

**Methods:** We present a statistical modeling framework to evaluate the spatial-temporal dynamics of the 2016-2017 dengue outbreak in the Negombo region of Sri Lanka as a function of human mobility, land-use, and climate patterns. The analysis was conducted at a 1 km × 1 km spatial resolution and a weekly temporal resolution.

**Results:** Our results indicate human mobility to be a stronger indicator for local outbreak clusters than land-use or climate variables. The minimum daily temperature was identified as the most influential climate variable on dengue cases in the region; while among the set of land-use patterns considered, urban areas were found to be most prone to dengue outbreak, followed by areas with stagnant water and then coastal areas. The results are shown to be robust across spatial resolutions.

**Conclusions:** Our study highlights the potential value of using travel data to target vector control within a region. In addition to illustrating the relative relationship between various potential risk factors for dengue outbreaks, the results of our study can be used to inform where and when new cases of dengue are likely to occur within a region, and thus help more effectively and innovatively, plan for disease surveillance and vector control.

## 1. Background

Dengue is a mosquito-borne viral disease that infects approximately 390 million people globally every year, particularly in tropical and subtropical countries (1, 2). The high number of infections combined with the lack, as yet, of a routinely used effective vaccine has made dengue a notorious public health problem (2, 3).

Dengue spreads through the bite of infected *Ades* mosquitoes, especially *Aedes aegypti*– the primary vector, with an estimated 15 to 17-day delay between the primary and secondary human infections (4). Dengue outbreak control is a challenge for policy makers because *Aedes aegypti* mosquitoes are well adapted to high density urban environments and actively feed during the day (5–7), thus presenting an elevated risk to humans. Urban settings provide an ideal habitat for *Aedes aegypti* breeding due to an abundance of discarded trash bags, plastic bottles, tires, and other containers that enable the formation of stagnant shallow water surfaces after precipitation (8). Urban regions in developing countries are particularly vulnerable due to a lack of indoor plumbing infrastructure that, in conjunction with a lack of air-conditioning, results in higher human-mosquito exposure rates during the day. Additionally, because of the daytime feeding behaviors of *Aedes aegypti*, common vector control measures that work for night-biting mosquitoes, such as bed nets, fail to effectively control dengue transmission. Given these challenges, there is a need to better understand and predict dengue outbreaks and transmission risk within urban regions in developing countries so that vector control and surveillance resources can be optimally allocated.

Previous studies highlighted human mobility as a critical factor for dengue transmission (9–15), which contrasts the more minor role travel plays in the spread of vector-borne diseases transmitted by night-biting mosquitoes (15). While *Aedes aegypti* mosquitoes have a hard time dispersing geographically across large areas because they rarely travel more than 400m from where they emerge as adults (16–19), humans regularly travel much longer distances on a daily basis. As new dengue cases and clusters are regularly reported kilometers apart, it is likely that human mobility play a critical role in the spread of dengue outbreaks, *i.e.*, infected humans introduce dengue into new mosquito populations at their trip ends. As an example, Vazquez-Prokopec, Montgomery (12) studied the pattern of dengue transmission using location-based contact tracing on infected dengue patients during a dengue outbreak centered at Cairns, Australia. They collected locations that the patients frequently traveled to during the daytime and 2-4 weeks prior to the onset of symptoms through phone interviews. The contact locations with a proximity of 100 meters and a separation of 20 days were spatial-temporally linked into pairs and then chains to identify the plausible sites of dengue virus transmission. They showed that the complex pattern of dengue transmission was primarily driven by human mobility, and that targeted residual spaying could potentially reduce the probability of dengue transmission up to 96%. Their study highlights the importance of understanding dengue transmission patterns to optimize the allocation of dengue prevention and vector-control measures.

In addition to human mobility, recent studies have pointed to a strong association between climate conditions and dengue outbreaks at various locations and across different temporal resolutions (8, 20–24). Precipitation, mean temperature and temperature fluctuation were revealed to affect the population dynamics of *Aedes aegypti* mosquitoes and the dengue virus extrinsic incubation period (25–29). Specifically, a suitable average temperature and moderate temperature fluctuations are often favorable for dengue transmission (25), while an increase in precipitation is strongly associated with the onset of a dengue outbreak (22). Humidity, a combined effect of precipitation and temperature, is also a common climate index to evaluate the environmental capacity for dengue emergence (20, 21, 30, 31). Wesolowski, Qureshi (13) accounted for both climate and mobility in a study of dengue virus transmission over a large dengue outbreak period in Pakistan. They developed an epidemiological model that included temperature and relative humidity as input parameters for mosquito dynamics, as well as biting rate to capture the interactions between human and mosquito hosts. Human mobility was captured using mobile phone data of ∼40 million subscribers to estimate the spatially explicit travel volume, albeit not differentiating infected and non-infected people. They showed that the emergence of dengue epidemics in a new region could be predicted using aggregated travel patterns from endemic areas in combination with the developed epidemiological model. While climatic factors were found to be significant for prediction, this was in part due to the large study region, *i.e*., country level, which has variable climatic suitability for the mosquito vector. The study region considered in our work is much smaller and has minimal climactic variability, thus alternative methods are required to distinguish site-specific risk.

Land-use patterns — indicators of human activities and potential breeding habitats — have also been linked to dengue outbreaks (32–37). Previous studies investigated the effect of land-use patterns on the spread of dengue and found that human settlements, water bodies, and mixed horticulture are the top three associated land-use patterns for dengue emergence in Malaysia (35). In another study (36), areas surrounded by rice paddies and marshes/swamps were associated with a significantly higher population of dengue vectors during the rainy season in Thailand. Orchards (which often contain artificial water containers) and irrigation fields have also been shown to play an important role in dengue infections; however, their role varies given different local conditions. Sometimes, land-use type can be a proxy for other features, such as socio-economic factors, which may have a contradictory effect on dengue infections (37).

In this study, we present a statistical modeling framework to evaluate the relative role of human travel patterns, climate conditions, and land-use patterns on dengue outbreak dynamics in Negombo, Sri Lanka (Figure 1), where dengue outbreaks are increasing in scale and prevalence. Dengue has been endemic in Sri Lanka since 1960s, and all four serotypes circulate in the region (38). Dengue hemorrhagic fever (DHF) cases were rare until 1989, when the first major dengue outbreak occurred with approximately 200 cases and 20 deaths; DHF has since become endemic in the country (38–40). The 2017 outbreak involved many severe DHF cases, with evidence showing a dengue virus type 2 (DENV-2) being the causative serotype (41, 42). DENV-2 had only be detected infrequently over the recent decades of dengue epidemics and is therefore associated with low immunity in the region (38). As a result, more than 80,000 dengue cases including 215 deaths were reported nationally in less than seven months between 2016-2017, a fourfold increase in the number of reported cases compared to the average number over 2010-2016 (43). The region of Negombo, located in the Western province, experienced the greatest number of dengue cases in the country; approximately 45% of the cases nationwide by July 2017 (Figure 1), and is the focus area of our study.

**Fig. 1.**
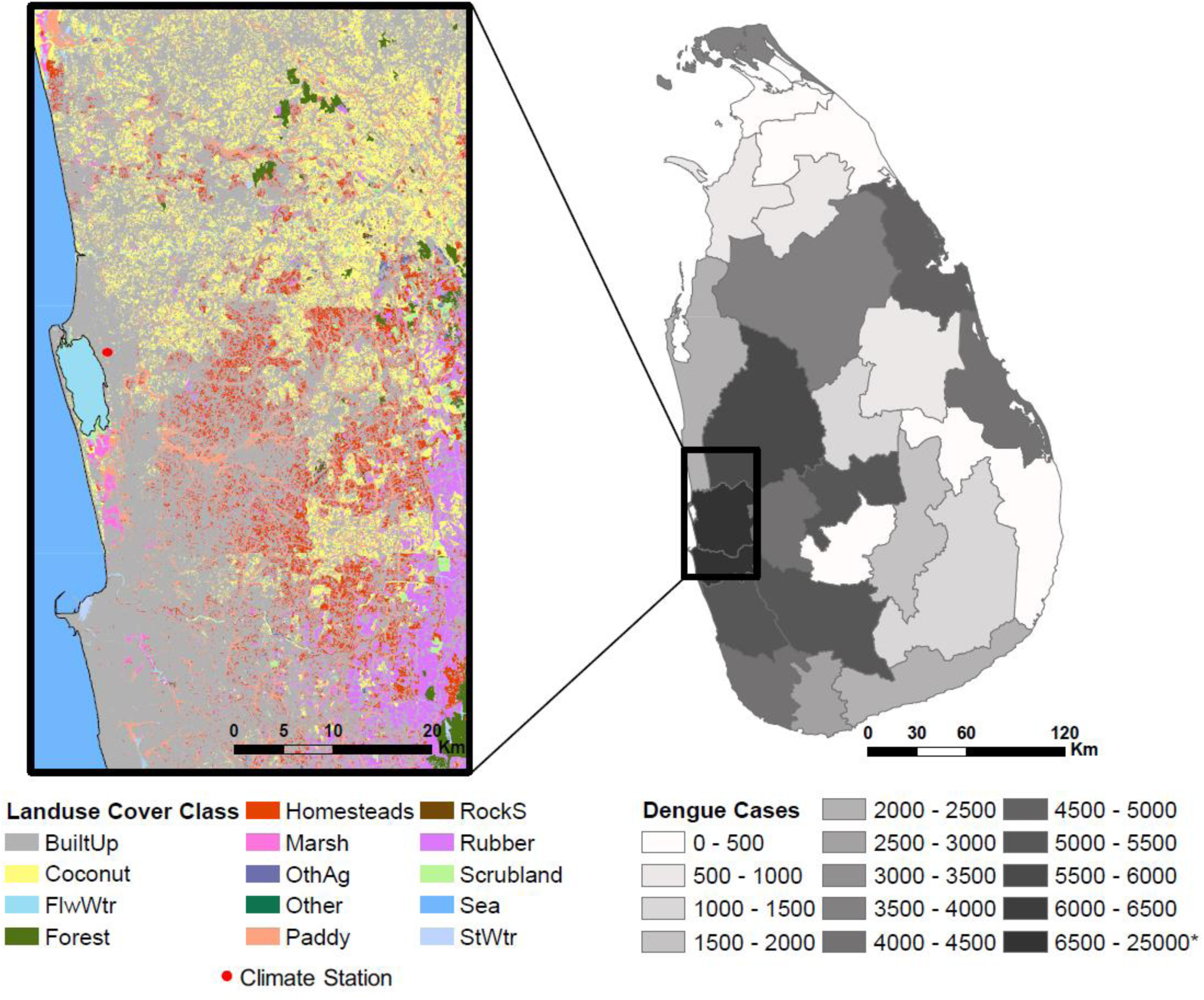
Land-use groups and the climate station in the study region of Negombo, Sri Lanka (left), and the total number of dengue cases at the district level in Sri Lanka (45) during the months from October 2016 to June 2017 that cover our study period (right).

We applied a mixed-effects model, where the mobility data bridges the time-varying, spatially-invariant climate variables and the rasterized spatially explicit, time-invariant population and land-use variables, to capture the spatial-temporal dynamics of dengue transmission. Our model framework differs from previous studies that simulated the transmission process (12, 13, 44), and instead focuses on estimating the timing and location of new case introductions though a non-process based statistical model. Specifically, we focus on modeling the home locations of (newly infected) dengue patients, and assume dengue is introduced in new areas by infected individuals who travel to the area. This assumption is consistent with previous studies that have shown visits to a household by infected people determines the infection risk in that household (11). In addition, the study was conducted at a fine-grained spatial and temporal resolution — 1 km × 1 km spatially and one week temporally — providing an improved understanding of the role of mobility in the spread of dengue. While previous work studied the impact of mobility (12, 13), climate (8, 20, 22, 25), and land-use (35–37) separately on dengue, the authors are unaware of any existing study that considers these factors within a single integrated framework. Thus, previous studies have been unable to quantify the relative contribution of each factor on the spatial-temporal patterns of dengue transmission as we do. The results from our study indicate that mobility is a more significant indicator of new dengue case clusters compared with land-use and climate factors. Furthermore, the case study in Sri Lanka provides critical insights into effective application of dengue prevention and vector control measures in developing regions.

## 2. Methods

A statistical model is applied to investigate the spatial-temporal dynamics of dengue outbreak with respect to a range of potential explanatory variables. The complete set of potential explanatory variables is listed in Table 1. Detailed descriptions of the data followed by a description of the methodology are provided below.

**Table 1:**
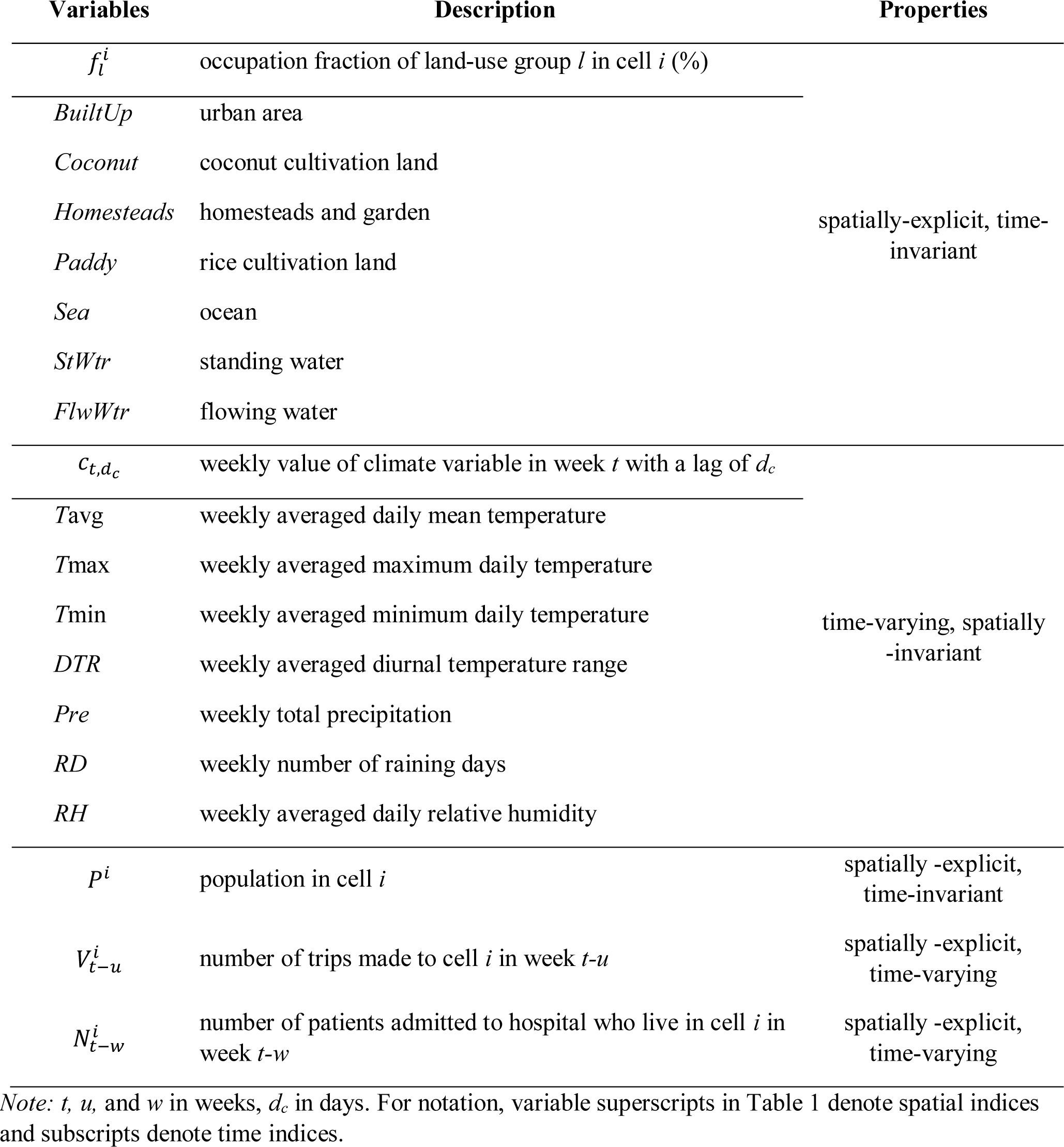
Summary of potential explanatory variables

### (a) Case and Mobility Data

A patient travel survey was conducted among dengue patients in the Negombo region of Sri Lanka over an approximately 8-month period during a major outbreak spanning from end of October 2016 to early July 2017. Geolocation data were collected from all patients admitted to the special High Dependency Unit (HDU) for critically ill dengue patients within the Clinical Centre for Managing Dengue and Dengue Haemorrhagic Fever (CCMDDHF) at the Negombo Hospital in Negombo, Sri Lanka. Specifically, the date of admission, home address, the complete set of locations visited, and corresponding trips made between all locations during the 10-days prior to hospital admission were collected from all HDU CCMDDHF admitted patients for the entire study period. The case data provide spatial-temporal information on the outbreak patterns, while the mobility data collected captures daily travel activity of the admitted dengue patients. The data were collected by trained students and supervised by a Senior House Officer on site. For weekday admissions the patients were surveyed upon admittance, for night admissions data were collected the following day, and for weekend admissions on the following Monday. The majority of admissions were 48 to 72 hours following the onset of fever. Dengue infection was confirmed for the patient set using either NS1 antigen or IgM antibody diagnostic test.

### (b) Climate Data

We used the Global Surface Summary of the Day (GSOD) daily weather data (46) from a station in Negombo (Figure 1) to explore the impact of climate factors on the dengue outbreak. The location of the weather station (7.18°N, 79.87°E) is approximately in the center of the study region and it is the only station that falls into our study region with a comprehensive set of climate data available during the study period. There are several global reanalysis products that provide spatial-explicit climate data during the study period; however, upon evaluation against the station observations, these globally gridded data sets did not provide accurate representations of the local climate variables, particularly at a daily time-step (Figure S1). Hence, the weather data are assumed to be representative for the region which has relatively homogeneous weather patterns (47). We selected a range of potential climate variables based on previous studies (8, 20–22, 25–31), including daily mean temperature (*T*avg), daily maximum temperature (*T*max), daily minimum temperature (*T*min), diurnal temperature range (*DTR*), precipitation (*Pre*), the number of raining days (*RD*), and relative humidity (*RH*) to analyze climatic influence for the weeks before and during the same period of analysis that the mobility data was collected.

### (c) Population and Land-use Data

We used a global population data layer based on Landscan 2016 (48), that is available at an approximately 1 km × 1 km resolution to represent the population distribution spatially. We aggregated the data to 5 km × 5 km grid for additional analysis with a coarser spatial resolution. Land-use data (49) were obtained from the Sri Lanka Survey Department which performed an initial survey in 2000 and has since continuously updated the maps. The map was extracted for our region of interest and reclassified into several groups (Figure 1): *Sea, Standing Water (StWtr), Flowing Water (FlwWtr), Coconut, Marsh, Paddy, Built-up (BuiltUp), Scrubland, Homesteads, Forest, Rubber, Rock/Sand (RockS), Other Agriculture (OthAg), and Other*. Water bodies were categorized depending on the potential effect on dengue transmission dynamics. Additional details on land-use classification groupings and processing is available in the supplementary material.

### (d) Data Processing and Statistical Model

We divided the study region (Figure 1) into a grid at a 1 km × 1 km resolution and aggregated daily data into a 1-weekly resolution. The number of patients who were admitted to the hospital during each week of the recorded time period was used to generate the weekly number of newly admitted dengue patients in each cell based on their home locations. This becomes our ‘case’ variable.

To incorporate the role of mobility into the model we used the travel itineraries provided by the patients to generate a time-dependent connectivity matrix, which represented the total number of trips made by dengue infected patients between each pair of cells for each week of the study period. The travel data included all destinations visited each day during the 10 days preceding hospital admittance (the time interval that the patient is assumed to be able to spread the disease) for each patient. The number of daily trips between each pair of cells was summed over all patients, to provide daily trip volumes between cells, and then aggregated to the weekly level. For each cell the total incoming weekly trips was summed to define our ‘trip’ variable. Critically, we exclude all trips with a destination of ‘home’ when computing our trip variable, in order to remove the inherent dependence between the ‘case’ variable, *i.e.*, the home location of infected individuals, and the ‘trip’ variable (explanatory variable). Thus, the total number of trips (excluding trips home) made by infected dengue patients entering a given cell *i* in a given week *t*, 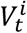, was used as a spatial-temporal explanatory variable in the model. The same method was used for the 5 km × 5 km analysis.

Climate variables were averaged or aggregated temporally to a weekly resolution, including weekly average *T*avg, *T*min, *T*max, *DTR, RH*, weekly total *Pre*, and *RD*. Land-use data were aggregated spatially to match the targeted spatial grid resolution. The population data were in an original resolution that matched the 1 km × 1 km grid. For land-use, the percentage of occupied land of each type was determined for each 1 km × 1km grid cell. Both were subsequently aggregated to a 5 km × 5 km grid.

A linear mixed-effects model combined with backward elimination of insignificant fixed effects (*p*-value > 0.05, two-tail test) was applied to investigate the spatial-temporal dynamics of dengue outbreak with the potential explanatory variables at a weekly time step and 1 km × 1 km spatial resolution. In building the model we first conducted sensitivity analysis to identify the optimal set of climatic variables to include in the model, and corresponding time lag for each of them.

Along with the chosen climate variable, the remaining set of potential explanatory variables (Table 1) was normalized and then taken into the mixed-effects model initially, with population included in the spatial random effects. Population density was incorporated using random effects in the model because population is likely to have spatially heterogeneous effects on dengue outbreaks (44, 50). For example, high population areas may imply access to tap water and better living conditions which could restrict dengue transmission (51), while the higher density of population facilitates disease spread. Furthermore, there could be spatial variance in the distribution of people living in a particular area. In addition to mobility, climate, and land-use variables; the number of new cases in a given cell in the weeks prior were added as explanatory variables to account for autocorrelations in the case data. Subsequently, the variable with the most insignificant fixed-effects coefficient was eliminated each iteration, until only variables with significant coefficients (at 95% significance level) remained in the model. A range of lead time for 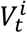 prior to the admitted week was also tested. A separate analogous process was conducted using a 5 km × 5 km resolution, to test the sensitivity of model results across spatial resolutions, and the robustness of the modeling framework and findings.

Thus, the mathematical representation of the model is given by:

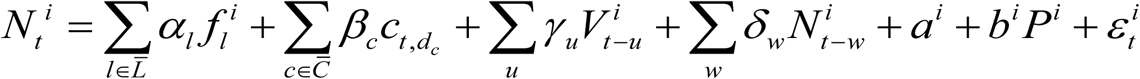

Where

*i* is the cell index; *i* = 1, 2, ….

*l* is the land-use variable, which belongs to the land-use group set 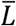, where 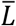 includes *Sea, StWtr, FlwWtr, Coconut, Marsh, Paddy, BuiltUp, Scrubland, Homesteads, Forest, Rubber, RockS, OthAg, and Other*.

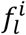 is the occupation fraction of land-use group *l* in cell *i*, time-invariant.

*p*^*i*^ is the population in cell *i*, time-invariant.

*t* is the time index at weekly resolution; *t* = 1, 2, ….

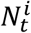 is the number of patients who are admitted to the hospital during week *t*, whose home locations are in cell *i*.

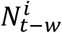 is the number of patients who are admitted to the hospital during week *t-w*, whose home locations are in cell *i*, where *w* is measured in weeks; *w* = 1, 2, …

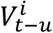 is the number of total number of trips made into cell *i* during the week *t-u*, where *u* is measured in weeks; *u* = 1, 2, …

*c* is the climate variable which belongs to the climate variable set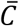. 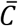 includes *T*avg, *T*max, *T*min, *DTR, Pre, RD*, and *RH*.

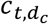 is the climate variable during the week that begins *d*_*c*_ days prior to the start of week *t*. *d*_*c*_ ranges from 7 to 17 days and can be different for different climate variables. Multiple climate variables can be included in the model.

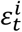 is the model residual associated with cell *i* and week *t*.

*α*_*l*_ is the estimated fixed-effects coefficient for *l*.

*β*_*c*_ is the estimated fixed-effects coefficient for *c*.

*γ*_*u*_ is the estimated fixed-effects coefficient for 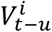.

*δ*_*w*_ is the estimated fixed-effects coefficient for 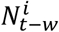.

*a*^*i*^ is the intercept associated with cell *i*.

*b*^*i*^ is the estimated spatial random-effects coefficient for *p*^*i*^.

The data processing and modeling were performed using MATLAB R2017a.

### (e) Sensitivity Analysis

We performed sensitivity analysis to evaluate the robustness of our model results. Firstly, a Jackknife analysis was conducted, specifically the statistical model was fit to all but one week of data, iteratively excluding one week at a time, over the entire time period modeled. The variability in estimated parameters and their corresponding significance are presented in Supplementary Table S3. Second, we evaluated the sensitivity of the model to fluctuations of the ‘trip’ variable. The motivation behind this sensitivity analysis was the uncertainty resulting from potential human error in the ‘trip’ variable, which is based on the patients’ recollection of their travel in the 10-day prior to hospital admittance. To assess the robustness of the model to error in the trip variable, we implemented Monte Carlo sampling, assuming an error of 10% uniformly distributed around the original observations 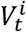. We performed 1000 random simulations, and the results are presented in Supplementary Table S4.

## 3. Results

### (a) Data Analysis

The number of admitted dengue patients aggregated over the study region peaks during December and June (Figure 2), aligned with the monsoon months (38). Figure 2a illustrates the relationship between the total number of dengue patients, *N*_*t*_, admitted during each week *t* and the total number of recorded patient trips (excluding the trips to home) during the same week (*V*_*t*_). Figure 2b illustrates *N*_*t*_ and the weekly averaged minimum daily temperature in week *t* (*T*min_*t*_). It shows a lagged relationship of *N*_*t*_ with *T*min_*t*_, mostly in the same direction. For the purposes of these graphics, the variables are aggregated over the entire study region.

**Fig. 2.**
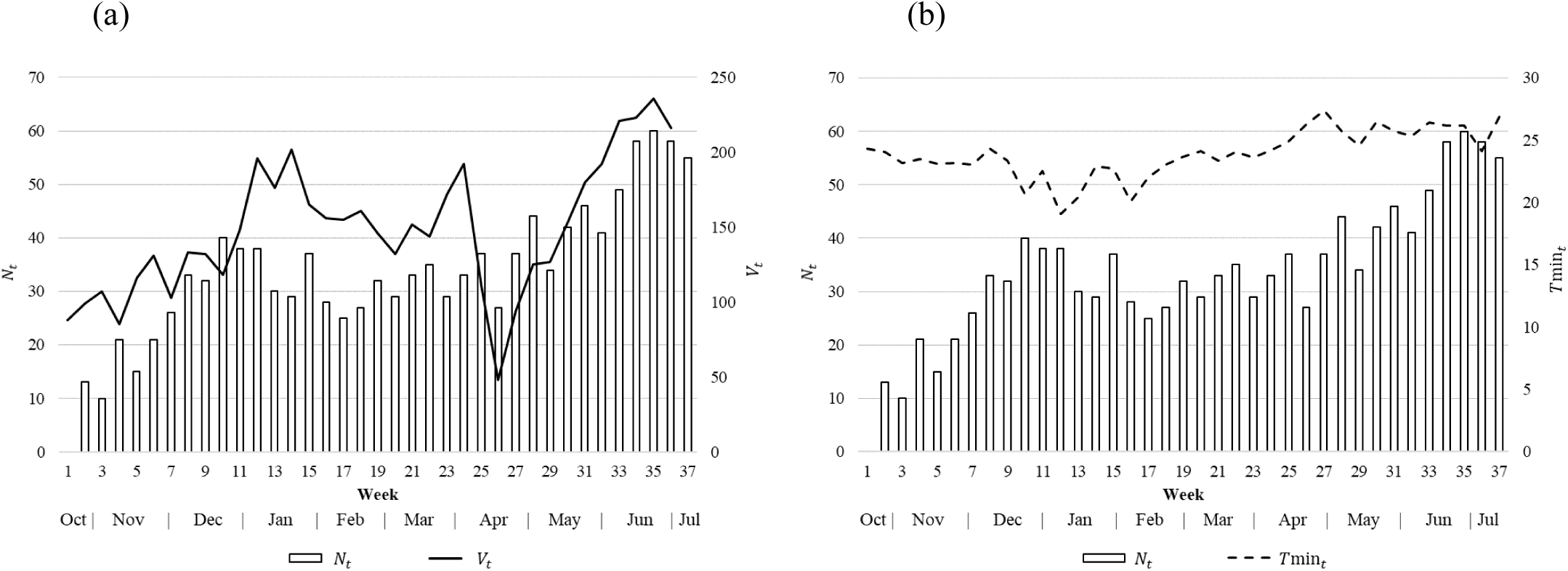
The number of admitted dengue patients in week *t* (*N*_*t*_) and (a) the number of recorded trips in week *t* (*V*_*t*_) summed over the entire study region, and (b) the weekly averaged minimum daily temperature (*T*min_*t*_).

The travel destinations recorded in our study include medical facilities, homes, workplaces, schools, and others (Figure S3). Additional analysis performed reveals that a vast majority of trips were longer than the distance a mosquito can travel. Specifically, 96.6% of the trips were longer than 0.4 km (Table S1; Figure S4), outside the range of a mosquito’s maximum travel distance (16–19), further supporting the role human mobility is likely to play in the outbreak.

Figure 3 illustrates both the spatial-temporal distribution of dengue patients’ home locations over the course of the outbreak, and the corresponding travel patterns of the patients during 5-week periods, excluding the trips with a destination of ‘home’. The patient home locations were well distributed over the area of the study region for the first few months of the outbreak, with correspondingly scattered travel patterns. However, as the outbreak progressed, the recorded case locations and the trip ends of newly infected dengue patients became more concentrated near the town center and just above the lake. There were also a large number of trips (>50) within the cell near the town center.

**Fig. 3.**
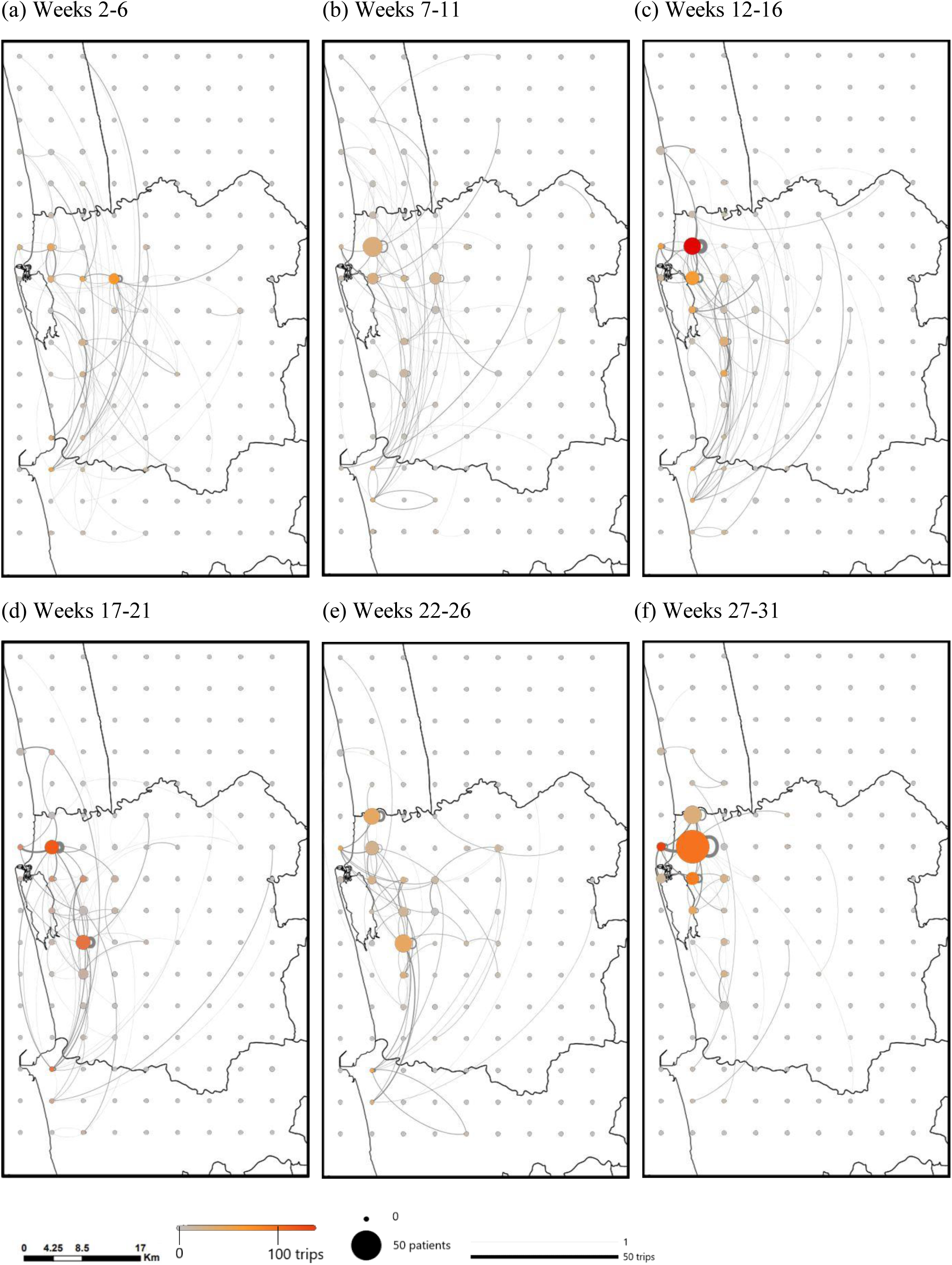
Weekly number of patients and the number of trips summed over 5-week intervals for a 5 km × 5 km resolution (excluding trips with a destination of ‘home’). Patient home locations are plotted as the case location. The size of the circle indicates the number of patients admitted during the time period. The color of the circle indicates the number of trips that end in the grid cell during the time period. The thickness of the line is proportional to the number of trips made between two locations. Week 2 begins on October 27, 2016 and week 31 begins on May 18, 2017. For visual clarity, the 5 km × 5 km resolution was used for the figure, instead of 1 km × 1 km.

### (b) Selection of Climate Variables

Among the climate variables, significant correlations were observed for weekly averaged *T*avg, *T*min, and *DTR* with a lead time ranging from 7 days to 17 days prior to the weekly admitted number of patients (*N*_*t*_), where the lead time (*d*_*c*_) is the lag in days between the climate variable and *N*_*t*_ (Figure S2). Regression models based on different combinations of the climate variables and lead time were developed and compared; the best performance model was select based on *F*-test and adjusted-*R*^2^. As a result, *T*min with an optimal lead time of 10 days was included in the final set of mixed-effects models to account for the partial influence of climate on the dengue outbreak (*R*^*2*^ = 0.248; *adj. R*^*2*^ = 0.226). This is consistent with previous findings (29) that daily minimum temperature were associated with increase in the larval abundance. We assumed a relatively homogenous climate over the study region, thus *Tmin* does not vary spatially over the study region.

### (c) Model Results

A mixed-effects model was developed to estimate the number of new dengue cases in a given cell in a given week as a function of the mobility patterns of individuals infected with dengue in the preceding week(s), as well as land-use and climate data from days prior.

Multiple models with explanatory variables representing land-use, climate, and mobility were created, and the three representative models are presented here. The three models vary based on the type of mobility variable included, specifically how far back in time travel is accounted for. The first model includes the mobility patterns one-week prior (*u* = 1), the second model includes the mobility patterns two-weeks prior (*u* = 2), and the third model excludes mobility altogether (“Exclude *V*”). The final set of climate and land-use variables found to be significant varies between models. All explanatory variables were normalized to a mean of zero and a standard deviation of one in the mixed-effects model. The fixed-effects coefficients (Table 2) therefore reflect the relative influence of each explanatory variable on the dengue outbreak dynamics.

**Table 2:**
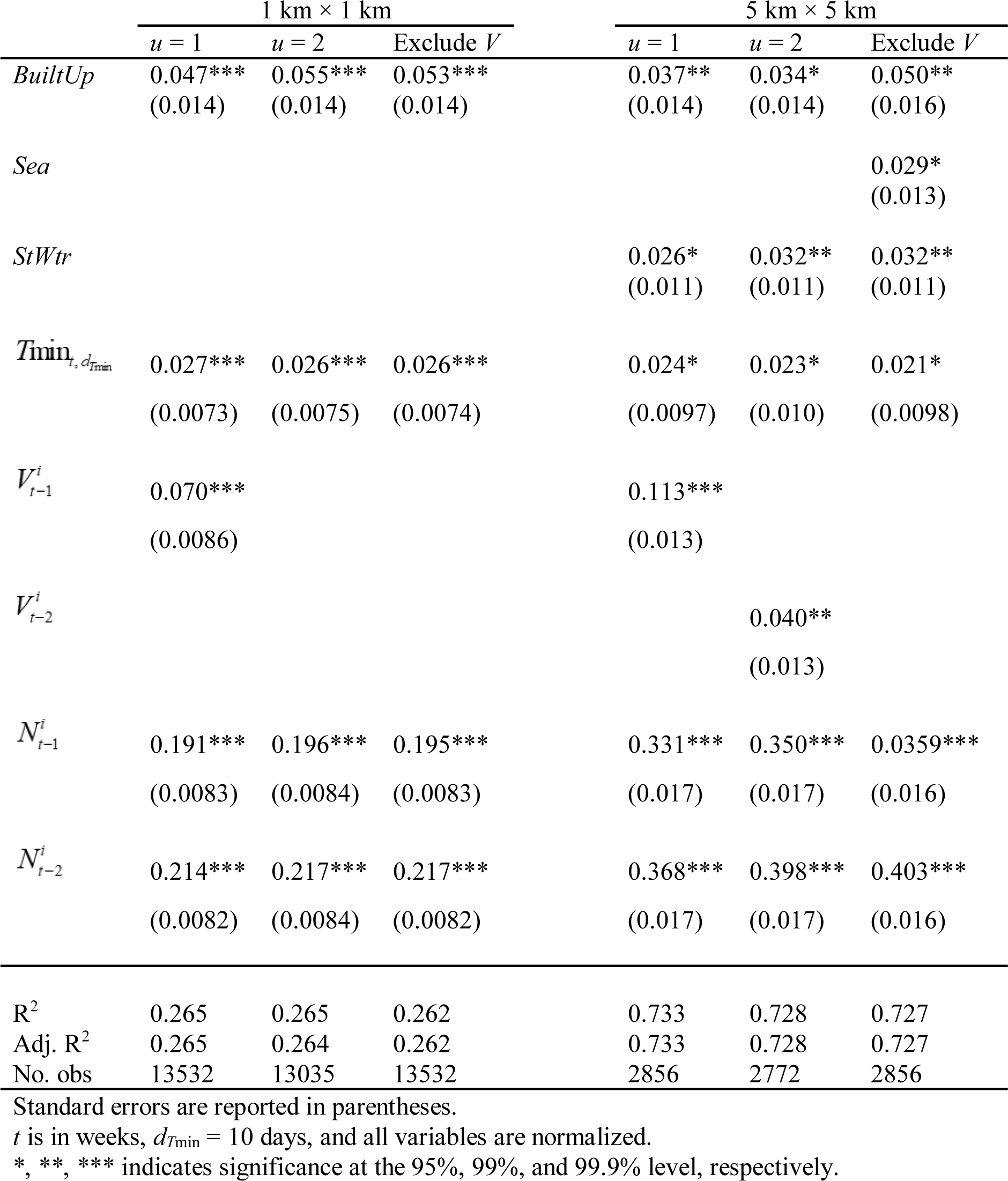
Fixed-effects coefficients and standard error of the mixed-effects model outputs based on the 1 km × 1 km and 5 km × 5 km resolution, respectively. The presented results are post-completion of the backward elimination of nonsignificant fixed effects. Variables without coefficients listed in the table were eliminated during the backwards elimination procedure for each model (each column). Variable descriptions are listed in Table 1.

The results (Table 2) for each of the three models are presented for both a 1 km × 1 km and 5 km × 5 km resolution, and reveal that the mobility patterns of dengue patients, specifically the number of trips made into a cell in a given week, to be the most dominant explanatory factor of new dengue cases in that cell the following week. Under the spatial resolution of 1 km × 1 km, the fixed-effects coefficient for the trips one week prior (*u* = 1) is 0.070, which is greater than the fixed-effects coefficients for other explanatory variables, suggesting human mobility plays a critical role in dengue outbreak dynamics. Results also illustrate a decrease in explanatory power of mobility patterns further than a week in advance, and are insignificant for the trips variable with a lead time of two weeks (*u* = 2). When mobility data is excluded from the model altogether, the adjusted *R*^*2*^ decreases slightly. In general, the power of the number of trips in predicting dengue cases deteriorates with longer lead time, with the number of trips two-weeks prior showing little or no advantage over other explanatory variables. The same conclusion is applicable for the results under the 5 km × 5 km resolution, as shown in Table 2. These results highlight the importance of collecting and utilizing mobility data within an appropriate timeframe for the purposes of modeling dengue outbreak risk. These conclusions are further supported by the sensitivity analysis, which show the rank and magnitude of the estimated model parameters (and significance) are robust to changes in both the outbreak period and also variability in the trips variable (Figure S6).

Among the seven land-use groups (see variable descriptions in Table 1) under the 1 km × 1 km resolution, only *BuiltUp* shows significant positive fixed effects on dengue cases (Table 2). Under the coarser spatial rasterization of 5 km × 5 km, *StWtr* and *Sea* also show significant positive fixed effects, in addition to *BuiltUp*. Whereas *BuiltUp* and *StWtr* show significant fixed effects in all three models with the coefficients ranging from 0.034 to 0.050 for *BuiltUp* and 0.026 to 0.032 for *StWtr*,, *Sea* shows the significant coefficient of 0.029 only in the model with the trip variable excluded. It indicates that urban areas, areas with standing water, and areas near the coastline are associated with a higher risk of dengue infections; the effect of *StWtr* and *Sea* is stronger under the 5 km × 5 km spatial resolution. In contrast, human mobility and *BuiltUp* are shown to be significant and robust indicators of dengue dynamics for both spatial resolutions. The same conclusions hold under the sensitivity analysis performed (Table S3 and S4).

Results from the sensitivity analysis are presented in Supplementary Table S3 for jackknife, and Table S4 and Figure S6 for trip variable error. Both sets of analysis reveal minimal fluctuation in the estimated model coefficients and set of significant variables, which illustrates the robustness of the model results. Specifically, for both spatial resolutions, travel volumes into a given cell one week prior is always a significant indicator of newly reported cases in the cell, while travel two-weeks prior is a less reliable factor, and negligent at smaller spatial scales, *i.e.*, 1 km × 1 km. Further, travel patterns one-week prior is consistently identified as a more significant indicator of new dengue cases than climate and land use.

## 4. Discussion

The results from this study illustrate the critical contribution of human mobility on the location and timing of new dengue cases, relative to land-use and climate variables. The results are sensitive to the temporal patterns of travel during the week immediately preceding the appearance of new case reports. This was the variable with the greatest predictive power. Although, travel patterns two weeks prior were found to have an insignificant effect on dengue outbreak dynamics. Our results are consistent with Stoddard, Forshey (11), who concluded that visits to households by dengue infected individuals determines the infection risk, further validating our use of patient home locations in the model. Furthermore, the significance of mobility in outbreak prediction was found to be robust under both spatial resolutions of 1 km × 1 km and 5 km × 5 km.

In contrast to the role of mobility, which we found to be a consistently significant indicator of new dengue cases, the effect of land-use patterns on the number of new cases is sensitive to the spatial resolution of the models. Land-use variables played a larger explanatory role at the coarser spatial resolution of 5 km × 5 km (compared to the finer 1 km × 1 km resolution), particularly for smaller spatially-dominant land-use patterns such as *Sea* and *StWtr*. *BuiltUp* showed the strongest positive effect overall, indicating urbanization is associated with an increased risk of dengue outbreak (which is consistent with multiple previous findings (52, 53)). *StWtr* also showed significant positive effect, which is to be expected because standing water provides suitable mosquito breeding habitat (19). The positive effect of *Sea* only appeared significant when human mobility was excluded from the model. Given the significant positive correlation between *Sea* and the number of trips (Table S2), it is likely that the large travel volume towards the area near the coastline makes the study region prone to dengue outbreaks. Previous studies have also found evidence that dengue mosquitoes can breed in brackish water (54), further supporting our finding of the distance to the sea as significant in some of the models. If this pattern holds in other regions, as seems likely, that fact can be used for the spatial prioritization of resource allocation for disease case and vector surveillance and control.

Among the climate factors, temperature-related variables including *T*avg, *T*min, and *DTR*, were more strongly associated with the outbreak emergence than precipitation-related variables including *Pre* and *RD*, or *RH*, which is related to both. This finding is in accordance with (22), which concluded that “rainfall strongly modulates the timing of dengue (*e.g.*, epidemics occurred earlier during rainy years) while temperature modulates the annual number of dengue fever cases.” Based on regression analysis, we found *T*min with a 10-day lead time to be the best climate-based predictor of new weekly dengue cases. Given the likely robustness of this result in other regions, this fact can be used for the temporal prioritization of resources.

In addition to human mobility, climate, and land-use variables, which were included as fixed effects, population density was incorporated using random effects in the model because population is likely to have spatially heterogeneous effects on dengue outbreaks, as noted in the *Methods*. Based on the model results, the random-effects coefficients for population are mostly positive, as expected, indicating that higher population density is associated with a higher number of dengue cases (Figure S5). The most significant positive effect is seen north of the lagoon along the coastline, highlighting potentially high-risk areas, where higher populations are likely to facilitate the emergence of dengue outbreaks. A few cells resulted in negative random-effects coefficients, which may be due to confounding interactions between different variables included in the model, or alternative factors not captured in the model; these cells were few and only occurred in the model when the dominant mobility variable was included. It is possible the dominant role of mobility could over compensate for the impact of population, *e.g.*, because people are likely to travel to crowded downtown areas, along the lagoon, or near the ocean where the large number of trips made to those regions could offset the impact of population. That the random-effects coefficients for population density are positive and negative lends support to the modeling decision to treat it as having enough stochasticity to qualify as a random-effects variable.

The results of this analysis have implications that are relevant to the design of measures to control dengue cases, such as allocation of resources for mosquito vector control. Previous global modeling of ecological suitability for dengue vector mosquito species (both *Aedes aegypti* and *Aedes albopictus*) have shown that the entire study area is a prime habitat for these species (55, 56). This conclusion drawn from the global models finds validation in our analysis, which shows that climate and land-use variables are not the most strongly associated with dengue case outbreaks. Consequently, epidemiological risk based on vector ecology may be insufficient for the purposes of optimizing vector control resource allocation, as it is unable to distinguish between potential sites to target within the study area. Because travel into the sites is the most important predictor of new case clusters, it may well be time to optimize vector control resources based on mobility data, with the aim to prevent exposure to the day-biting mosquitoes, *i.e., Aedes aegypti*, at the highest risk locations. To the best of our knowledge, such a design for dengue control measures has not yet been tried in the field.

Finally, various limitations of this study should be noted. First, only dengue patients admitted to the CCMDDHF at the Negombo Hospital were included and surveyed in this study. Thus mild or asymptomatic cases, which account for the majority of dengue cases (1), were not accounted for in the study. Second, some patients infected in the study region may have gone to hospitals in other districts and would therefore not be included. Third, the mobility data was based on patients’ recollections over a 10-day period prior to hospital admission, and therefore may have inaccuracies due to human error in recalling the information. However, detailed analysis of the travel data revealed most trips recorded represent daily commuting routines. Thus, while some trips may be excluded due to human error, we believe the relative connectivity between cells is accurately captured by the survey responses. In addition, the sensitivity analysis revealed our results are robust to variation in the time of outbreak and trips variable. Fourth, the distance traveled and the time spent in a certain location were not considered due to the unavailability of relevant data; however these factors have been shown to have little influence on dengue transmission (11), and thus their exclusion does not invalidate the methodology used in this analysis. Fifth, by utilizing all the mobility data collected, we made an implicit assumption that the patients were infectious during the entire 10-day period prior to hospital admittance. This period does fall within the combined intrinsic incubation period (4-10 days) (57) and the early symptomatic period before admitted to the hospital. A sixth assumption was that the patients were infected at or around their home locations. This assumption is consistent with a wide variety of previous studies that revealed homes as the primary point of contact for dengue transmission (11, 58, 59). Vazquez-Prokopec, Montgomery (12) tried to identify the most plausible transmission locations based on reported contact locations from a dengue outbreak in Cairns, Australia and found that only 10.2% of the identified transmission sites were at out-of-home locations, and a notable portion of them were within 1 km of the home locations. Given that our objective was not to model the transmission chains of dengue as in (12), assuming home locations as the site of infection provides reasonable support for predicting where infected individuals reside, and therefore the risk posed around homes of infected individuals. Lastly, the climate data were obtained from a single station, thus a homogenous climatic region was assumed for our study region. Therefore, the role of climate factors on the dengue outbreak may be underestimated.

While the modeling framework used here is readily applicable to other contexts, future work should investigate how widely transferable the model results are. More specifically using general mobility data (tracking movements for all residents); such as using mobile phone data as in (13), or transport planning data, which may be more readily available and cost effective; should be compared to the use of patient mobility surveys as in this study.

## 5. Conclusions

Our study highlights the potential value of using travel data to target vector control within a region. In addition to illustrating the relative relationship between various potential risk factors for dengue outbreaks, the results of our study can be used to inform where and when new cases of dengue are likely to occur within a region, and thus help more effectively and innovatively, plan for disease surveillance and vector control.

## List of abbreviations

DENV-2: dengue virus type 2
HDU: High Dependency Unit
CCMDDHF: Clinical Centre for Managing Dengue and Dengue Haemorrhagic Fever
GSOD: Global Surface Summary of the Day
Tavg: daily mean temperature
Tmax: daily maximum temperature
Tmin: daily minimum temperature
DTR: diurnal temperature range
Pre: precipitation
RD: the number of raining days
RH: relative humidity
StWtr: Standing Water
FlwWtr: Flowing Water
BuiltUp: Built-up
RockS: Rock/Sand
OthAg: Other Agriculture

## Declarations

### Ethics approval and consent to participate

The protocol was reviewed and approved by the Johns Hopkins Bloomberg School of Public Health Institutional Review Board (JHSPH IRB), #IRB00009485, as well as approved by the Negombo Hospital. Patient consent was obtained in writing from all participants for the purposes of this study.

### Concent for publichation

Not applicable

### Availability of data and material

Patient case reports and travel diaries can not be shared due to privacy restrictions. The remaining data used in this study is provided as a supplementary file, along with the code used to generate the results.

### Authors’ contributions

L.G. conceived the study. Y.Z., S.Siddique and L.G. designed the experiments. Y.Z., H.S., L.F. and J.R. collected the data. Y.Z. developed the model and performed the computational analysis. All authors analyzed the data and model results. Y.Z., S.Siddiqui., S.Sarkar, J.R., K.O., and L.G., contributed to the writing of the manuscript.

## Acknowledgements

We thank Dhammika Silva from the Centre for Clinical Management of Dengue and Dengue Haemorrhagic Fever who supervised the data collection and Benjamin Zaitchik at Johns Hopkins University for advice on the climate data extraction.

