## Supplementary figures and tables for "Modeling the relative role of human mobility, land-use and climate factors on dengue outbreak emergence in Sri Lanka"

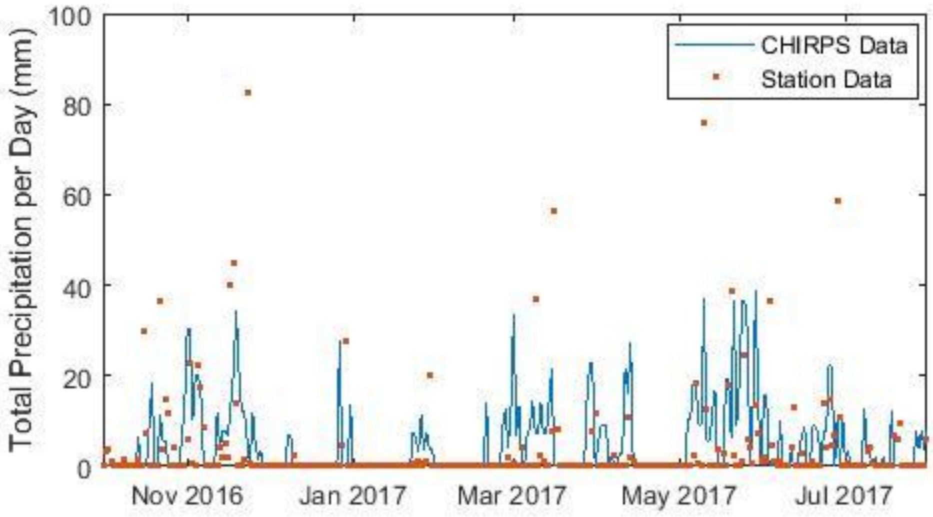


**Fig. S1.** Daily rainfall time series from the climate station used in the study and the global precipitation reanalysis product Climate Hazards Group InfraRed Precipitation with Station data (CHIRPS).

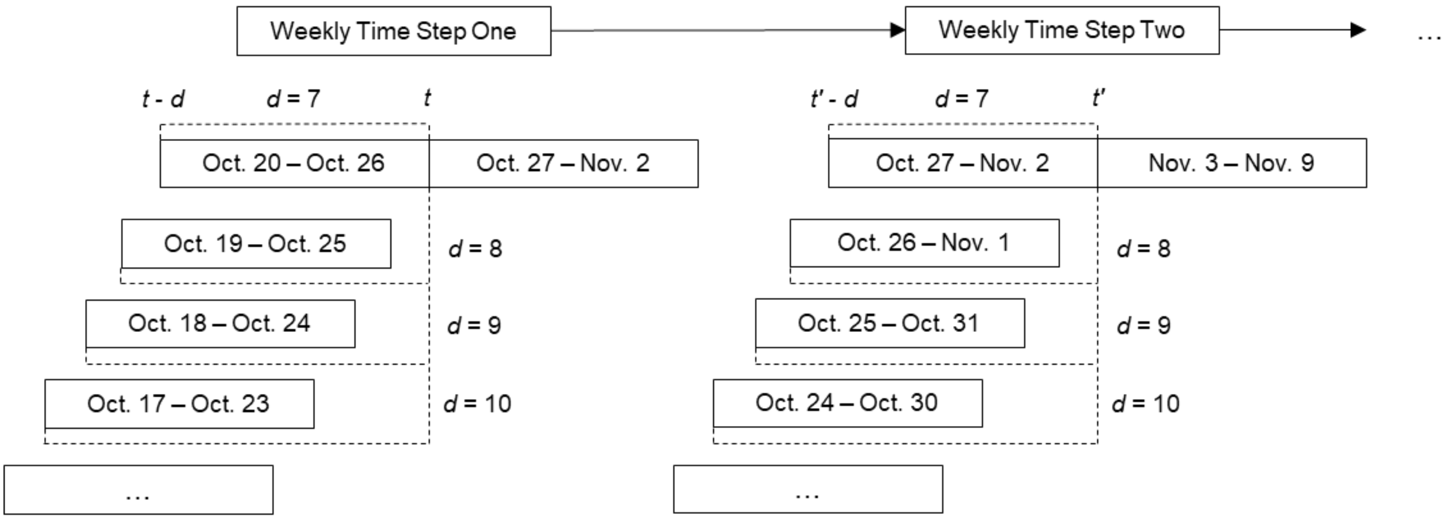


**Fig. S2.** Illustration of the definition of lead time *d* in days within a temporal framework of weekly time steps *t*. As illustrated, *t* - *d* is the week that begins *d* days prior to the start of week *t*. A range of lead time *d* from *d* = 8 to *d* = 10 prior to the week *t* with calendar dates listed are shown as examples. As a result, weekly averaged *Tmin* with a lead time of 10 days (*d* = 10) was included in the model (*Results*).

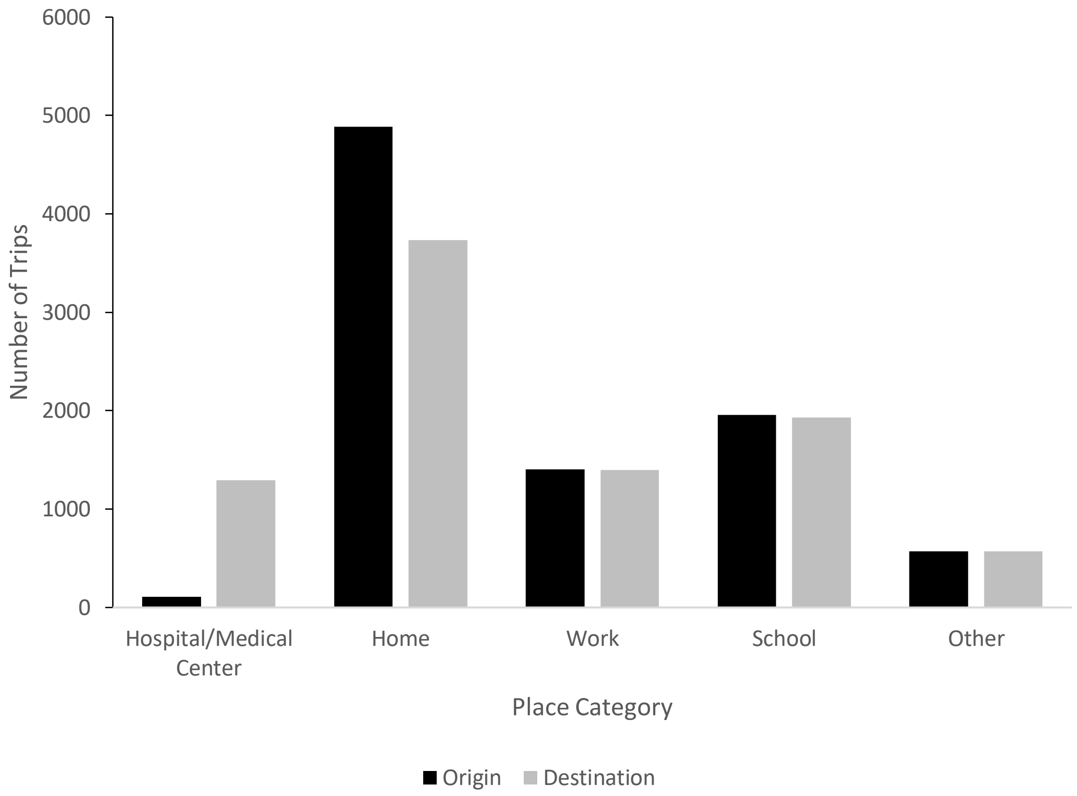


**Fig. S3.** Origins and destinations of trips categorized by trip end location.

**Table S1: Number of trips Categorized by length**

| Trip Length (km) | Frequency |
|---|---|
| 0-0.4 | 252 |
| 0.4+ | 7377 |

*Note*: trips to hospitals were excluded

(A) (B)

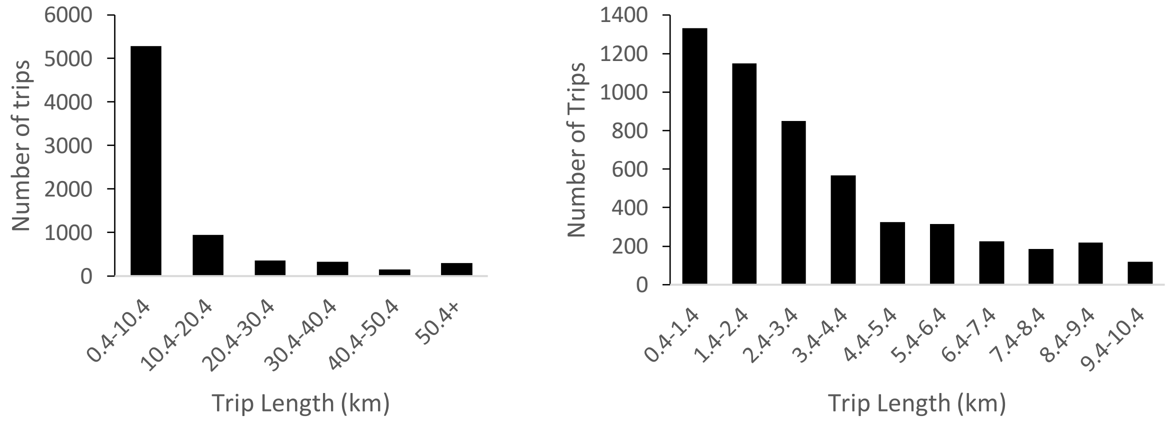


**Fig. S4.** Distribution of trip lengths for trips over 0.4 km. (A) All trips above 0.4 km. (B) Trips between 0.4-10.4 km. Note trips with medical facilities as the trip destination were excluded.

**Table S2: Correlations between explanatory variables**

| | | $N_t^i$ | *BuiltUp* | *Sea* | *StWtr* | $T\mathrm{min}_{t,\, d_{T\mathrm{min}}}$ | $V_{t-1}^i$ | $V_{t-2}^i$ | $N_{t-1}^i$ | $N_{t-2}^i$ |
|---|---|---|---|---|---|---|---|---|---|---|
| | | | | | | 1 km × 1 km | | | | |
| $N_t^i$ | | | 0.08*** | 0.00 | 0.00 | 0.04*** | 0.54*** | 0.31*** | 0.38*** | 0.39*** |
| *BuiltUp* | | 0.17*** | | -0.22*** | -0.16** | 0.00 | 0.13*** | 0.13*** | 0.08*** | 0.07*** |
| *Sea* | | 0.00 | -0.40*** | | -0.02 | 0.00 | 0.02* | 0.02* | 0.00 | 0.00 |
| *StWtr* | | 0.13*** | -0.17 | 0.10 | | 0.00 | 0.02* | 0.01 | 0.00 | -0.01 |
| $T\mathrm{min}_{t,\, d_{T\mathrm{min}}}$ | 5 km × 5 km | 0.05** | 0.00 | 0.00 | 0.00 | | 0.00 | -0.01 | 0.03*** | 0.02** |
| $V_{t-1}^i$ | | 0.82*** | 0.16*** | 0.09*** | 0.17*** | 0.00 | | 0.64*** | 0.37*** | 0.30*** |
| $V_{t-2}^i$ | | 0.71*** | 0.17*** | 0.09*** | 0.17*** | -0.01 | 0.86*** | | 0.54*** | 0.36*** |
| $N_{t-1}^i$ | | 0.79*** | 0.16*** | 0.00 | 0.13*** | 0.05* | 0.75*** | 0.80*** | | 0.36*** |
| $N_{t-2}^i$ | | 0.80*** | 0.16*** | -0.01 | 0.13*** | 0.03 | 0.74*** | 0.74*** | 0.77*** | |

*, **, *** indicates significance at the 95%, 99%, and 99.9% level, respectively.

**Table S3: Jackknife sensitivity analysis.** The average fixed-effects coefficients and Jackknife standard error of the mixed-effects model outputs are shown for the 1 km × 1 km and 5 km × 5 km resolution. The presented results are post-completion of the backward elimination of nonsignificant fixed effects. Variables without coefficients listed in the table were eliminated during the backwards elimination procedure for each model (each column). Variable descriptions are listed in Table 1.

| | 1 km × 1 km | | | 5 km × 5 km | | |
|---|---|---|---|---|---|---|
| | *u* = 1 | *u* = 2 | Exclude *V* | *u* = 1 | *u* = 2 | Exclude *V* |
| *BuiltUp* | 0.047*** | 0.055*** | 0.053*** | 0.038** | 0.036 | 0.051*** |
| | (0.0085) | (0.0098) | (0.0084) | (0.011) | (0.024) | (0.0095) |
| *Sea* | | | | | | 0.030** |
| | | | | | | (0.0092) |
| *StWtr* | | | | 0.027* | 0.032*** | 0.033*** |
| | | | | (0.012) | (0.0061) | (0.0053) |
| $Tmin_{t,\, d_{Tmin}}$ | 0.027*** | 0.027*** | 0.027*** | 0.025*** | 0.023*** | 0.022*** |
| | (0.0041) | (0.0045) | (0.0043) | (0.0067) | (0.0060) | (0.0054) |
| $V^i_{t-1}$ | 0.071** | | | 0.117* | | |
| | (0.020) | | | (0.046) | | |
| $V^i_{t-2}$ | | | | | 0.040 | |
| | | | | | (0.051) | |
| $N^i_{t-1}$ | 0.193*** | 0.198*** | 0.197*** | 0.337*** | 0.355*** | 0.364*** |
| | (0.041) | (0.042) | (0.040) | (0.065) | (0.060) | (0.059) |
| $N^i_{t-2}$ | 0.210*** | 0.213*** | 0.213*** | 0.363*** | 0.390*** | 0.396*** |
| | (0.056) | (0.058) | (0.056) | (0.096) | (0.090) | (0.087) |
| $R^2$ | 0.263 | 0.263 | 0.260 | 0.731 | 0.726 | 0.725 |
| Adj. $R^2$ | 0.263 | 0.263 | 0.260 | 0.730 | 0.726 | 0.725 |
| No. obs | 13019 | 12527 | 13019 | 2756 | 2672 | 2756 |

Jackknife standard errors are reported in parentheses.
*t* is in weeks, $d_{Tmin}$ = 10 days, and all variables are normalized.
*, **, *** indicates significance at the 95%, 99%, and 99.9% level, respectively.

**Table S4: Sensitivity analysis of 'trip' variable.** The average fixed-effects coefficients and standard errors are based on 1000 random simulations. For the variables that are not significant in every simulation the percentage indicates how often it occurred as significant. The coefficient values are averaged over the significant outputs.

| | 1 km × 1 km | | 5 km × 5 km | |
|---|---|---|---|---|
| | *u* = 1 | *u* = 2 | *u* = 1 | *u* = 2 |
| *BuiltUp* | 0.048*** | 0.055*** | 0.037** | 0.035* |
| | (0.014) | (0.014) | (0.014) | (0.015) |
| *Sea* | | | | 0.029* |
| | | | | (0.013) |
| | | | | 4.9% |
| *StWtr* | | | 0.026* | 0.032** |
| | | | (0.011) | (0.011) |
| $Tmin_{t, d_{Tmin}}$ | 0.027*** | 0.027*** | 0.024* | 0.022* |
| | (0.0073) | (0.0075) | (0.0097) | (0.010) |
| $V^i_{t-1}$ | 0.061*** | | 0.109*** | |
| | (0.0085) | | (0.013) | |
| $V^i_{t-2}$ | | 0.018* | | 0.038** |
| | | (0.0086) | | (0.013) |
| | | 3.0% | | 95.1% |
| $N^i_{t-1}$ | 0.192*** | 0.196*** | 0.332*** | 0.351*** |
| | (0.0083) | (0.0084) | (0.017) | (0.017) |
| $N^i_{t-2}$ | 0.214*** | 0.217*** | 0.370*** | 0.399*** |
| | (0.0082) | (0.0084) | (0.017) | (0.017) |
| $R^2$ | 0.265 | 0.265 | 0.733 | 0.728 |
| Adj. $R^2$ | 0.264 | 0.264 | 0.732 | 0.728 |
| No. obs | 13532 | 13035 | 2856 | 2772 |

Standard errors are reported in parentheses.
*t* is in weeks, $d_{Tmin}$ = 10 days, and all variables are normalized.
*, **, *** indicates significance at the 95%, 99%, and 99.9% level, respectively.

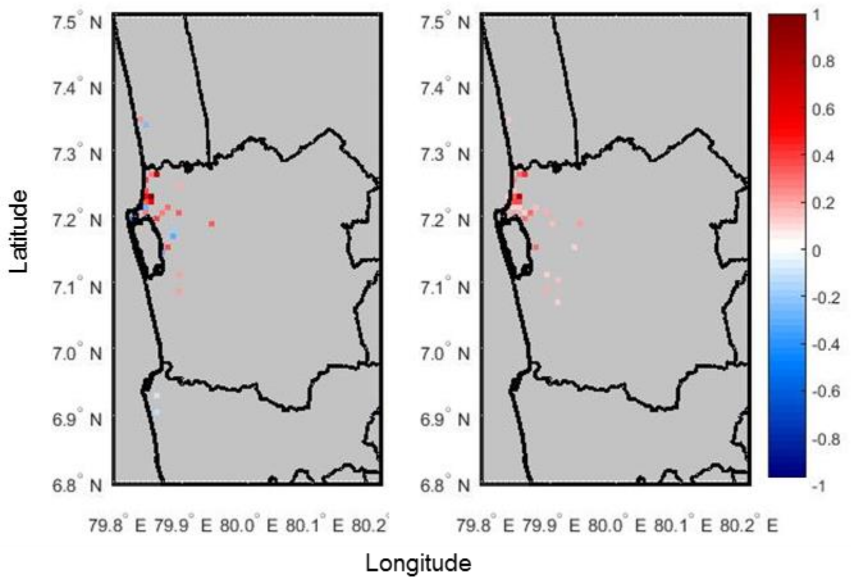


**Fig. S5.** Random-effects coefficients for population on a color scale from -1 to 1 for model with the number of trips one week prior (left) and with the trip variable excluded (right), under 1 km × 1 km spatial resolution. Only significant coefficients at 95% level are shown (nonsignificant coefficients are marked in grey).

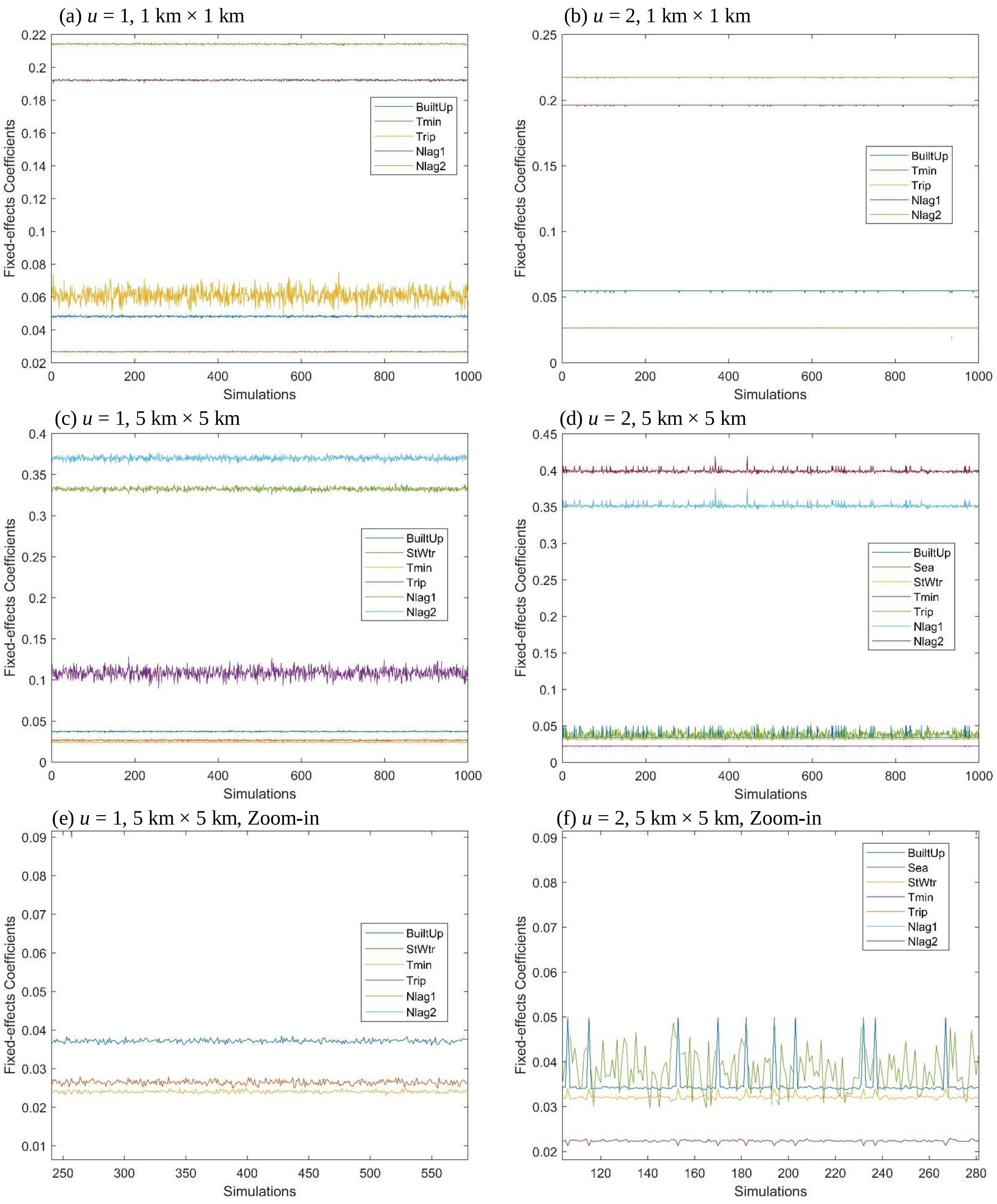


**Fig. S6.** Significant fixed-effects coefficients of explanatory variables under 1000 random simulations for the sensitivity analysis of ‘trip’ variable. Each colored line represents the fixed-effects coefficients that showed up significant for an explanatory variable over the 1000 simulations given different models with (a) $u = 1$, *i.e.*, mobility patterns one-week prior, and for a spatial resolution of 1 km × 1 km, (b) $u = 2$ for 1 km × 1 km, (c) $u = 1$ for 5 km × 5 km, and (d) u = 2 for 5 km × 5 km. (e) and (f) are zoom-in figures of (c) and (d), respectively. From all figures, one can see the ranking details among the lines, *i.e.*, the comparison of the magnitude of the fixed-effects of different explanatory variables under the random simulations for the sensitivity analysis.

### Land-use Data

Dengue vector mosquitoes are known to reproduce in areas where standing water such as puddles can form after rain events. Therefore, a cover class of standing water had to be distinguished from flowing water bodies or ocean. Standing water included abandoned irrigation channels and tanks (reservoirs), ponds, and lakes. Ocean was also distinguished from flowing water class because high salinity conditions do not enable the reproduction of mosquitoes.

Vegetation cover was separated into agricultural and non-agricultural areas. Agricultural areas were further categorized into *Coconut*, *Paddy*, *Rubber and OthAg*. Because these areas generally consist of monocultures, differences in fertilizer, insecticide or other chemical use can potentially play a significant role in the lifecycle and reproductive cycle of mosquitoes. Similarly, differences in conditions necessary to grow these vegetation types can also affect mosquito populations. *Paddies* were distinguished due to the presence of standing water necessary for the cultivation of rice. Nonagricultural areas consisted of *Homesteads*, *Scrubland*, *Marsh*, *Forest*, and *RockS*.

The *BuiltUp* cover contained developed areas – generally locations with building structures and a high percentage of impervious surface, representing a large portion of cities and towns. The original map obtained from the Sri Lanka Survey Department, however, classified the city of Negombo and certain parts of Colombo as *Coconut*. Using Landsat-8 images courtesy of the U.S. Geological Survey (1), a supervised classification was performed in ArcMap 10.4.1 to more precisely delineate the extent of the *BuiltUp* layer. Specific areas on the Landsat-8 image that were known to be built-up were highlighted and marked as such using the classification tool. Their aggregated spectral signature was extracted and subsequently used as a reference marker for the *BuiltUp* cover class. Areas that were not built-up were aggregated together and their spectral signature was extracted. Within the supervised classification tool, the spectral signature data were run through a maximum likelihood algorithm that allowed the tool to select which pixels were more likely than not to be classified as being *BuiltUp*. These pixels were used as the new *BuiltUp* extent and replaced previous cover class in the map.
